# Do host-consumed resources increase endoparasitic but decrease ectoparasitic infections?

**DOI:** 10.1101/2021.05.12.443795

**Authors:** Brin Shayhorn, Chloe Ramsay, Kristi Medina, Erin Sauer, Jason R. Rohr

**Author notes:** Co-first authorship (Brin Shayhorn, Chloe Ramsay). Corresponding Author: Chloe Ramsay, Department of Biological Sciences, 100 Galvin Life Science Center, University of Notre Dame, IN, 46556.

## Abstract

Habitat loss and disease outbreak play a major role in the decline of biodiversity. Habitat degradation is often associated with reduced food resources, which can lead to less investment in host immunity and increased infections. However, pathogens use host resources for replication and pathogen traits, such as infecting hosts internally or short generation times, might allow pathogens to rapidly capitalize on host-consumed nutrients. Thus, it is unclear whether increased food consumption by hosts should reduce or amplify pathogen levels. We conducted experiments on Cuban treefrogs (*Osteopilus septentrionalis*) to test how food availability affects infection levels of Ranavirus and the fungal pathogen *Batrachochytrium dendrobatidis* (Bd), which are both associated with mass die-offs of amphibians. Given that Ranavirus is an endoparasite with a much shorter generation time than the ectoparasitic Bd, we postulated that Ranavirus might be able to capitalize on host-consumed resources more quickly than Bd. We hypothesized that increased food availability to hosts might reduce Bd infections more than Ranavirus infections. As predicted, augmenting food access decreased Bd infection intensity, but increased Ranavirus infection intensity. Future work should assess whether pathogen traits, such as generation time and endo- versus ectoparasitism, generally affect whether food resources more positively benefit hosts or pathogens.

## Introduction

Habitat loss and degradation can reduce food resources for wildlife, which can exacerbate and cause disease (Daszak et al. 1999, Gibbons et al. 2000). This is because mounting an immune response requires resources and therefore hosts short on resources are unlikely to mount a completely effective immune response, which could increase disease risk and severity (Gray et al. 2009, Echaubard et al. 2010). As an example, food restricted bumblebees (*Bombus terrestris*) experienced a 50% increase in mortality from *Crithidia bombi* infection (protist) when compared to bees fed *ad libitum* (Brown et al. 2000).

While hosts use resources to combat pathogens, pathogens also utilize host resources. Thus, positive relationships between host resources and pathogen loads are theoretically possible (Cressler et al. 2014a). For example, increased resources can increase pathogen density in the environment and consequently pathogen load in hosts (Johnson et al. 2007, Civitello et al. 2015, Civitello et al. 2018). Additionally, hosts with more resources are often healthier and larger and thus might be more capable of tolerating higher infection loads (Pulkkinen and Ebert 2004, Knutie et al. 2017). Finally, high resource levels can be used by some pathogens to increase their replication, despite the potential for a subsequent increase in host immune response (Johnson et al. 2007, Seppälä et al. 2008, Hall et al. 2009, Cressler et al. 2014a, Civitello et al. 2018).

Although there have been in-depth examinations of how host traits and resources affect whether the host or parasite benefits more from increased resource availability (Pulkkinen and Ebert 2004, de Roode et al. 2008, Seppälä et al. 2008, Hall et al. 2009, Cressler et al. 2014b, a, Hite and Cressler 2019), surprisingly, the role of parasite traits on resource-dependent parasitism remains under tested. Certain parasite traits may facilitate pathogens utilizing host resources. For example, quickly replicating pathogens could potentially capitalize on high resource environments faster than slowly replicating pathogens (Arsnoe et al. 2011, Hall et al. 2012), and perhaps even faster than the host can use these resources to mount an immune response. Also, pathogens with more direct access to host-consumed resources, such as endoparasites in the gastrointestinal tract of the host, may be able to access these resources more easily than those without direct access, such as non-blood-feeding ectoparasites.

In recent decades, amphibian populations have experienced global declines, partially due to land use changes that have altered resource availability and disease outbreaks (Gibbons et al. 2000). Two pathogens contributing to population declines in amphibians are *Batrachochytrium dendrobatidis* (Bd) and Ranavirus (Daszak et al. 1999). Bd is a widespread and pathogenic fungus that attacks amphibian epithelial tissues, preventing osmotic regulation through the skin and causing mortality by cardiac arrest (Voyles et al. 2009). It is an ectoparasite, residing on the skin of amphibians with a ~ 4d replication rate at room temperature (Piotrowski et al. 2004, Voyles et al. 2009). Ranavirus is a pathogen that results in hemorrhaging and cell death of major organs (Gray et al. 2009). Ranavirus resides internally and has a comparatively high replication rate, inducing host death in as few as two days for tadpoles and seven days for adults of susceptible species (Green et al. 2002, Sutton et al. 2014).

Studies examining how host susceptibility to Bd and Ranavirus are affected by resource availability have reported mixed results. One study found that increased resources reduced host susceptibility to Bd (Venesky et al. 2012) and the rest found no effects of resource availability (Reeve et al. 2013, Cothran et al. 2015, Buck et al. 2016). However, all these Bd studies focused on nitrogen, phosphorus, carotenoids, or protein levels, addressing food quality rather than quantity. Also, all these studies were conducted on tadpoles rather than adults, despite adults being the most susceptible amphibian life stage to Bd (Venesky et al. 2012, Cothran et al. 2015, McMahon and Rohr 2015, Buck et al. 2016). A study on how resource availability affected Ranaviral infections in tadpoles, the life stage when hosts are most susceptible to Ranavirus (Green et al. 2002, Gray et al. 2009), found that resources did not affect host susceptibility to Ranavirus (Reeve et al. 2013). However, this study only limited host access to resources before exposing hosts to Ranavirus (Reeve et al. 2013). Here, we examine how host-consumed resources affect Ranavirus and Bd growth on amphibians to gather evidence in support of or against the hypothesis that rapidly replicating endoparasites (i.e., Ranavirus) and slowly replicating ectoparasites (i.e., Bd) benefit more than and less than hosts, respectively, from host-consumed nutrients.

## Methods

### Animal Husbandry

Adult and larval Cuban tree frogs (*Osteopilus septentrionalis*) were collected from the University of South Florida Botanical Gardens (Tampa, Florida, USA) in November 2015 and August 2017, respectively. Both life stages were housed in 946 mL sterilized plastic deli cups, with adults placed on unbleached paper towels wet with artificial spring water (ASW) and larvae placed in 800 mL of ASW. All individuals were kept at 20°C with a 12hr photoperiod. Adults and tadpoles were used in Bd and Ranavirus experiments, respectively. While tadpole and adult frogs have physiological differences that change how they respond to pathogens, these life stages are the most susceptible to these pathogens (Gray et al. 2009, McMahon and Rohr 2015). Therefore, understanding resource use in these life stages is more ecologically pertinent than testing resource use at the same life stage for both pathogens.

### Experimental Design

*Bd experiment:* Eighteen adult Cuban tree frogs of equal mass were assigned randomly to the four treatments (2×2 experiment) for a total of 72 individually housed frogs. All individuals then received 1 mL of deionized water pipetted on to their dorsal surface that was from rinsed Petri dishes (15 x 150 mm; 1% agar, 1% tryptone medium) that either were (Bd treatment; *n* = 36) or were not (control; *n* = 36) growing Bd (SRS 812 isolate). Each Bd exposed frog received 10^5^ zoospores (Gervasi et al. 2013). After Bd or sham exposure, individuals were placed on either an *ad libitum* diet, where they were provided with an excess of small house crickets (*Acheta domestica*; ~0.27g) three times per week or a restricted diet, where they were provided ~0.094g of crickets three times per week.

*Ranavirus experiment:* Ten Cuban tree frog tadpoles were assigned randomly to the six treatments groups (2 x 3 experiment) for a total of sixty tadpoles individually housed in deli cups with 100mL of ASW. Ranavirus (FV3-like) was cultured in fathead minnow (*Pimephales promelas*) cells and maintained at −80°C in minimal essential medium (MEM). Tadpoles received either 77μL (10^5^ PFUs) of Ranavirus (FV3-like) in MEM or 77μL of MEM-without virus as a sham treatment. Previous studies found that doses between 10^2^-10^6^ PFUs cause sublethal effects and morbidity in tadpoles (Hoverman et al. 2010). After twenty-four hours, 700mL of ASW was added to each cup for a total of 800 mL/tadpole. Each tadpole was then provided either one, two, or three 0.6 x 1-cm cylindrical plugs (measured using the first centimeter of a disposable micropipette tip) of a mix of fish flakes and spirulina (1:1) in 1% agar, according to their randomly assigned low, medium, or high food treatment.

### General Methods

All amphibians were weighed or staged (if tadpoles; Gosner 1960) at the beginning of the experiment and every week thereafter. Clean containers were provided weekly. To quantify infections, each adult frog was swabbed weekly on their abdomen and hind limbs. The mouth and cloaca of tadpoles were swabbed at the end of the experiment. All swabs were placed in 1mL microcentrifuge tubes and stored at −20°C for later processing. After the experiment concluded, DNA was extracted from the swabs and quantitative polymerase chain reaction (qPCR) was used to quantify Bd load (Boyle et al. 2004) and Ranavirus load (Picco et al. 2007). Any individuals surviving to the end of the experiment (three weeks for Bd and two weeks for Ranavirus) were euthanized with a 0.1% solution of buffered MS-222.

For the Bd experiment, blood from euthanized frogs was extracted using microcapillary tubes to collect plasma. We then conducted enzyme-linked immunosorbent assays (ELISA) on the plasma to quantify IgY antibody levels (adapted from Knutie et al. 2017; see Supplemental Methods for details). We chose to focus on IgY levels because IgY is the most common immunoglobin and used to combat Bd (Grogan et al. 2018).

### Statistical Analysis

All analyses were conducted in R (3.4.2, 2017) and figures were created using the *visreg* package and *visreg* function (Breheny and Burchett 2019). We used general linear models to examine the effect of resource availability, parasite exposure, and their interaction on host growth, development (Gosner stage; Ranavirus experiment only), and immune response (IgY antibody levels in the Bd experiment only). Host growth was measured as the difference between pre-exposure and 3-week post-exposure body mass (g) for Bd and as change in mass per week for Ranavirus.

We also examined the effect of resource availability and Ranavirus exposure on survival by conducting a survival analysis using the *survival* package and the *coxph* function (Therneau and Lumley 2019). Individuals that survived to the end of the experiment were right-censored. No amphibians died in the Bd experiment and consequently survival analyses were not run.

To assess how Bd loads changed over time with resource availability, we analyzed Bd infection intensity using a generalized linear mixed effects model with a negative binomial error distribution. Resource availability and weeks since the beginning of infection were used as interacting independent variables, and individual was included as a random effect. To assess how Ranavirus intensity changed with resource availability, we used a similar analysis, but without the inclusion of week since infection or the random effect in the model because intensity was not measured through time. To assess how resource level affected pathogen prevalence, we ran two generalized linear regression models (one for each pathogen) with binomial error distributions. The binomial response variable was host infection status (infected or not) and resource level was used as a predictor variable. Generalized linear mixed models were run using the *pscl* package and *glm.nb* function (Jackman et al. 2020). Tukey post-hoc tests were run to test which resource levels differed from one another using the *multcomp* package and *glht* function (Hothorn 2010).

## Results

In both the Bd and Ranavirus experiments, hosts fed more had higher growth rates (g/week; Ranavirus: *F* _2, 51_=5.72, *p*<0.006, Fig. 1A; Bd: β=0.208, □^2^_1_= 58.196, *p* < 0.001, Fig. 1B). Although hosts at different life stages were fed different food types, the positive association between resource levels and growth rates across life stages suggests that the chosen food levels likely forced immunity/growth tradeoffs at the lower, but not at the higher resource levels across both life stages. However, neither Ranavirus exposure (β= −2.97, *F*_1, 51_=0.016, *p*=0.89) nor Bd exposure (β=−0.013, □^2^_1_= 0.478, *p* = 0.490) affected host growth rate.

**Figure 1.**
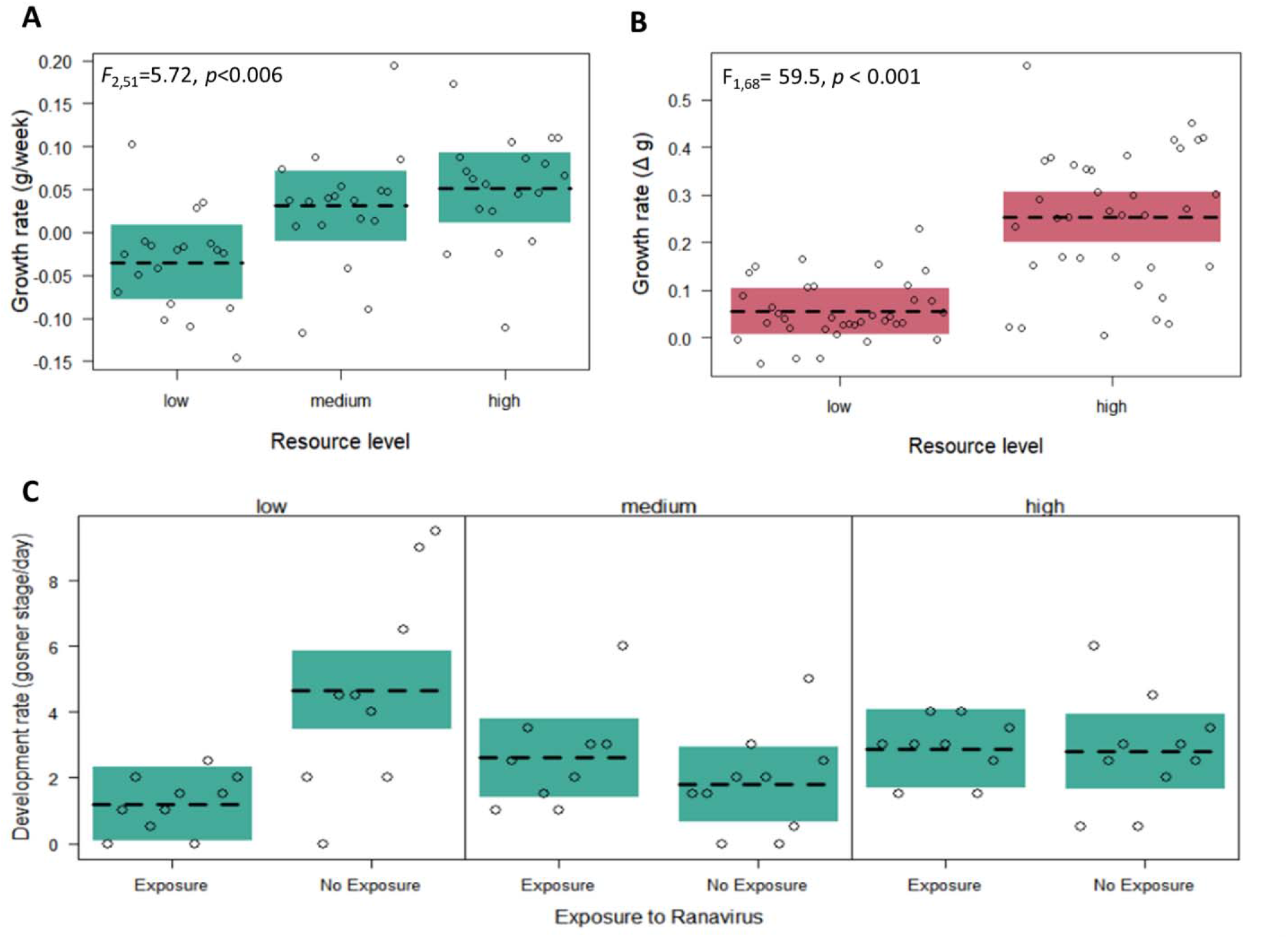
Effects of varying food levels on Cuban tree frog (*Osteopilus septentrionalis*) weight change (grams/week) when **A)** exposed or not to Ranavirus or **B)** *Bactrachochytrium dendrobatidis* (Bd) and **C)** on their development rate (Gosner stage/week) when exposed to Ranavirus or not. Increased food availability increased host growth rates (Ranavirus: *F*_2,51_=5.72, *p*<0.006; Bd: F_1,68_= 59.5, *p* < 0.001), regardless of pathogen exposure, whereas Ranavirus exposure significantly decreased development rates in the low resource treatment, but not across the other tested resource levels (resource x exposure: *F*_2,51_=7.89, *p*=0.001; low resource exposure: *F*_1,17_=10.92, *p*=0.004). Data are shown as a conditional plot, with expected values (line), 95% confidence intervals for the expected values (shading), and partial residuals (points).

Host development rate, as measured by Gosner stages per week, was significantly affected by an interaction between Ranavirus exposure and resource availability (*F*_2,51_=7.89, *p*=0.001). Ranavirus exposure decreased host development rate for hosts in the low resource group (*F*_1,17_=10.92, *p*=0.004), but did not significantly affect development rate in the medium or high resource groups (medium resources: *F*_1,17_=1.32, *p*= 0.27; high resources: *F*_1,17_=0.02, *p*=0.89; Fig. 1C).

Mortality was not significantly affected by resource availability (χ^2^_2_=0.0003, *p*=0.999), Ranavirus exposure (β=2.028, χ^2^_1_=0.33, *p*=0.567), or the interaction between the two (χ^2^_2_=3.94, *p*=0.14) in the Ranavirus experiment. One individual died in the control-low resource treatment group and one in the Ranavirus-low and Ranavirus-high resource treatment groups for a total of three mortalities. There was no mortality in the Bd experiment. Ranavirus intensity was 4.6x higher in tadpoles assigned to the high resource than the low resource group (χ^2^_1_=4.50, *p*=0.03; Fig 2A). In contrast, Bd intensities were 2.5x higher in frogs fed the low resource level than the high resource level (χ^2^_1_=4.27, *p*<0.05; Fig 2A), but there were no significant effects of time since infection (χ^2^_1_=079, *p*=0.78) or an interaction between resource availability and time (χ^2^_1_=0.51, *p*=0.47). Surviving tadpoles provided the low resource treatment had 89% Ranavirus prevalence, whereas tadpoles in the medium and high resource groups had 70% and 44% Ranavirus prevalence, respectively (χ^2^_1_=3.57, *p*=0.059; Fig. 2B). Bd prevalence did not significantly differ between the high (88%) and low (80%) resource groups (χ^2^_1_=0.50, *p*=0.48). Additionally, host IgY was not affected by resource availability (β=-0.12, *F*_1,32_= 2.34, *p* = 0.14), Bd exposure (β=0.059, *F*_1,32_= 0.083, *p* = 0.78), or their interaction (*F*_1,32_= 0.20, *p* = 0.66).

**Figure 2.**
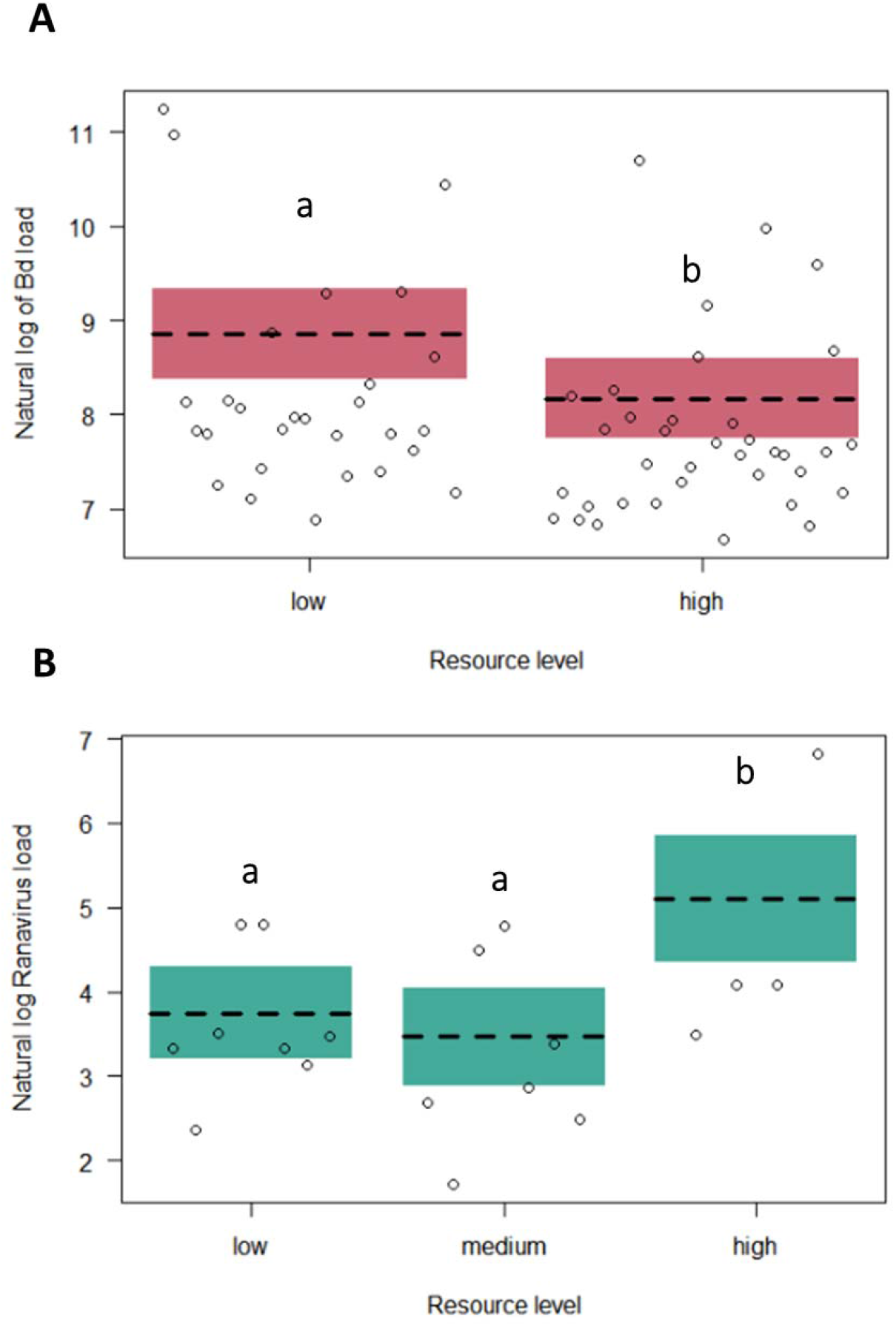
Pathogen intensity in infected larval (A) and adult (B) Cuban tree-frogs (*Osteopilus septentrionalis*) fed a range of resource levels (high, medium, and low). **A)** *Bactrachochytrium dendrobatidis* (Bd) loads were consistently higher in individuals with low resource availability (χ^2^_1_=4.27, *p*<0.05), which was independent of weeks since infection (time since infection-by-resource availability: χ^2^_1_=0.51, *p*=0.47). **B)** Ranavirus loads were higher when hosts were fed more food (χ^2^_1_=5.17, *p*<0.05). Data in both figures are shown as partial residual plots, with the expected values (line), 95% confidence bands or intervals for the expected values (shading), and partial residuals (points).

## Discussion

Hosts use resources to combat infections and limited resources may force tradeoffs between immunity and growth or reproduction (Lochmiller and Deerenberg 2000, Rauw 2012). Pathogens also use host resources making it unclear whether host-consumed resources should be beneficial or costly to the pathogen. Resource quantity or quality and host traits might determine whether increased resources are more beneficial for the host or pathogen (Cressler et al. 2014a). In this experiment, we assessed the level of evidence in support of the hypothesis that pathogen traits can alter competition for resources between host and pathogen. We found that resource level was positively associated with pathogen load for Ranavirus, a rapidly replicating endoparasite, but negatively associated with pathogen load for Bd, a more slowly replicating ectoparasite (Fig. 2).

In both experiments, increasing food availability increased frog growth, regardless of whether the frogs were exposed to pathogens or not (Fig. 1A, B), suggesting that hosts in the high resource groups had sufficient resources to combat pathogens. However, neither Ranavirus nor Bd infections affected host growth rate (g/week) or survival, regardless of resource availability. In contrast to growth rates, when hosts were exposed to Ranavirus and had limited resources, hosts had significantly decreased development rates, whereas Ranavirus exposure did not alter development rates when hosts were in the medium or high resource treatment groups (Fig. 1C). Amphibian larvae might increase development to escape a resource-limited environment, as has been seen with other inhospitable conditions, such as with predators (Relyea 2007), pesticides (Rohr et al. 2004), and in rapidly drying freshwater environments (Bekhet et al. 2014).

While higher resource levels did not alter IgY levels, they did decrease Bd intensity in hosts, suggesting that these increases in resources allowed hosts to more effectively combat Bd (Fig. 2A). Increased resources could have led to increases in immunity other than IgY, such as increases in antimicrobial peptides used to combat Bd, or increases in non-immunological defenses, such as increased skin shedding rates, which have been shown to decrease Bd intensity (Rollins-Smith et al. 2002, Cramp et al. 2014). Our results suggest that any increases in Bd replication from increased resources was not sufficient to counteract subsequent increases in immunity, leading to the highest Bd loads when the host had the least food. Bd is an ectoparasite and has a ~4 day replication rate, which could make it challenging for Bd to use host-consumed resources effectively (Piotrowski et al. 2004). This pattern of increased food resources decreasing Bd load was shown in a past study on larval southern leopard frogs (Venesky et al. 2012). However, other studies found that the quality of resources do not alter Bd loads in three different species of larval frogs (Cothran et al. 2015, Buck et al. 2016). Unlike these previous studies, our study tested resource quantity and not quality, which could account for these differences. Additionally, amphibian immunity in the larval stage is less complex than in the adult life stage; thus, how hosts use resources to combat Bd in the larval life stage might be different than in the adult life stage that we tested (Rollins-Smith 1998).

In contrast to Bd, abundant food increased Ranaviral intensity (Fig. 2B) while decreasing Ranavirus prevalence. Ranavirus grows best in hematopoietic tissues (Chinchar 2002), suggesting that Ranavirus may benefit from the increased growth we found in hosts with access to more resources. Also, Ranavirus replicates rapidly and is an endoparasite, which may allow this pathogen to more effectively use host resources for replication than some more slowly replicating ectoparasites, like Bd. In a past study on wood frogs (*Rana sylvatica*), reduced resources did not affect host susceptibility to Ranavirus (Reeve et al. 2013). However, in this study frogs had reduced access to resources only before individuals were infected, which means that Ranavirus likely had similar levels of host-consumed resources to utilize (Reeve et al. 2013). In our study, Ranavirus prevalence decreased with increasing resources, the opposite of the intensity pattern. These results together suggest that increased resources may help the host mount an initial immune response to combat Ranaviral infections, but once established, Ranavirus can use host consumed resources to rapidly replicate.

Our experiments did not test the same host life stage or resource type, and thus we cannot rule out these differences as explanations for our findings. We chose to study different host life stages because we wanted to test particularly harmful (i.e., deadly) pathogens during the host life stage when they are most susceptible (Gray et al. 2009, McMahon and Rohr 2015). Otherwise, differences among the pathogens in their responses might simply be due to low growth rates and thus low statistical variance on the host life stage where the pathogen performs most poorly. Importantly, Bd has much lower growth rates and is less virulent in tadpoles than postmetamorphic frogs, not because of superior immunity at this stage, but because of a lack of substantial keratin, the resource for the pathogen. In fact, although there are differences between the immune systems of tadpoles and adults, both life stages have functioning immune responses to combat and clear the tested pathogens (Rollins-Smith 1998, Gantress et al. 2003, Grayfer et al. 2012, Fites 2014). Previous work has demonstrated that protein quantity in food is an important predictor in how hosts respond to infection (Sandland and Minchella 2003, Venesky et al. 2012). However, both spirulina (fed to the tadpoles) and crickets (fed to the adult frogs) have similarly high protein levels (60-65%; NOW Spirulina, Bloomingdale, IL, USA, Mariod et al. 2017), suggesting that protein differences are unlikely to account for our results. Despite not being able to rule out life stage or diet differences as explanations for our results, we believe that the most parsimonious explanation is that the observed patterns reflect differences in host and pathogen resource use. We encourage future studies to test the hypothesis that rapidly replicating endoparasites and slowly replicating ectoparasites benefit more than and less than hosts, respectively, from host-consumed nutrients using a single host life stage and virulent pathogens.

Organisms in natural ecosystems often experience resource limitation from natural or anthropogenic factors, driving the importance of understanding how resources alter disease progression in hosts. In our study, higher resources increased amphibian growth but altered pathogen abundance positively or negatively depending on the pathogen. These results support the hypothesis that pathogen traits may be an important factor in determining whether the host or pathogen benefit more from increased resource availability. Other studies have found that resource availability is positively associated with pathogen replication in viruses (Arsnoe et al. 2011, Hall et al. 2012) and in endoparasites (Pulkkinen and Ebert 2004, Tylianakis et al. 2004), consistent with our results and the notion that rapidly replicating endoparasites might benefit from increases in host-consumed resources. Future studies should further examine the role that pathogen traits play in determining whether the host or pathogen benefit more from increased resource availability.

## Acknowledgements

We would like to thank the University of South Florida for allowing access to use their Botanical Gardens for specimen collection. Thanks also to Jason Hoverman who supplied the Ranavirus, Joyce Longcore who supplied the Bd used for infecting amphibians, and Sarah Knutie for developing and assisting with the ELISA assay used in this experiment. JRR was supported by grants from the National Institutes of Health (R01TW010286-01), and the National Science Foundation (DEB-2017785, IOS-1754868).

## Authorship Contributions

All authors conceived ideas and designed methodology. BS and KM collected the data. CR analyzed the data. BS and CR led the writing of the manuscript. All authors contributed critically to the drafts and gave final approval for publication.

## Data Availability

Upon publication the data will be made available on Dryad.

