## Supplemental Methods for "Do host-consumed resources increase endoparasitic but decrease ectoparasitic infections?"

We then conducted enzyme-linked immunosorbent assays (ELISA) on the plasma to quantify IgY antibody levels (adapted from Knutie et al. 2017). Only adult amphibians from the Bd experiment were used this for immune assay because adult amphibians have a more complex and well-developed immune response than tadpoles (Rollins-Smith 1998, Robert and Ohta 2009). Serum from individual frogs was first diluted at 1:100 in carbonate coating buffer (0.05 M, pH 9.60) and then added to ninety-six well plates with 100 μL/well. Each individual sample was run in triplicate. After serum was added, plates were incubated at 4 °C overnight. Plates were washed five times with 300 μL/well of a Tris-buffered saline wash solution. The plates were then filled with 200 μL/well of bovine serum albumin (BSA) blocking buffer and incubated for 2 h at room temperature on an orbital table. Plates were washed again five times, loaded with 100 μL/well of primary detection antibody (Goat-αAlligator-IgG, diluted 1:1000; Bethyl), placed on an orbital table for 1 h, washed again (five times), loaded with 100 μL/well of secondary detection antibody (Rabbit-αGoat-IgG, diluted 1:5000; Bethyl) for 1 h, and then washed for the final time (five times). Plates were then loaded with 100 μL/well of tetramethylbenzidine (TMB: Bethyl Laboratories) and incubated for 30 mins before the reaction was stopped with 100 μL/well of stop solution (Bethyl Laboratories). Finally, optical density (OD) of the wells was measured with a spectrophotometer (BioTek, PowerWave HT, 450-nanometer filter).
